# Expansion of CRISPR loci with multiple memories of infection enables the survival of structured bacterial communities

**DOI:** 10.1101/747212

**Authors:** Nora C. Pyenson, Luciano A. Marraffini

## Abstract

Type II CRISPR-Cas systems provide immunity against phages and plasmids that infect bacteria. Following infection, a short sequence of the phage genome known as the “spacer” is inserted into the CRISPR locus to capture a memory of the infection and immunize the host. Spacers are transcribed and processed into guide RNAs that direct the Cas9 nuclease to its target on the invader. Thousands of spacers are acquired to target the viral genome at multiple locations and neutralize phage mutants that evade the immunity specified by a single guide RNA. In liquid cultures, where phages and their hosts are constantly mixed, spacer diversity is generated at the population level, and a single immunization per cell is sufficient to confer robust immunity. Although rare, bacteria that acquire multiple spacers can also be found, demonstrating that type II CRISPR-Cas systems also have the capability of generating spacer diversity at the cellular level. However, conditions in which this feature is important for survival are poorly understood. Here we found that when phage infections occur on solid media, a high proportion of the surviving colonies display sectored morphologies that contain individual cells with multiple spacers. We show that this is the result of the bacteria-host co-evolution, in which the immunity provided by the initial acquired spacer is easily overcome by escaper phages that decimate all the progeny of the founder cell that do not acquire additional spacers. Our results reveal the versatility of type II CRISPR-Cas immunity, which can respond with both single or multiple spacer acquisition schemes to solve challenges presented by different environments.

## Introduction

Clustered, regularly interspaced, short, palindromic repeats (CRISPR) loci and their associated (*cas*) genes provide immunity against viruses [1] and plasmids [2] that infect prokaryotes. Phage- and plasmid-derived “spacer” sequences that are intercalated [3–5] between repeats provide the specificity of the CRISPR-Cas immune response. This is because spacers are transcribed and processed into small RNAs (known as CRISPR RNAs, crRNAs) that guide Cas nucleases [6] to destroy complementary nucleic acids of the invaders [7–11]. New spacers are acquired at a low rate from the invader’s genome upon infection and, remarkably, invariably inserted in the first position of the CRISPR array [1]. Therefore, spacer acquisition is comparable to an immunization of the bacterial or archaeal host and the CRISPR array provides a temporal “vaccination” record of the immunization events.

Depending on the *cas* gene content of a CRISPR locus, CRISPR systems are classified into six different types (I-VI) [12]. Here we investigated spacer acquisition by the *Streptococcus pyogenes* type II-A CRISPR-Cas system, which employs the RNA-guided nuclease Cas9 to provide immunity through cleavage of the invader’s DNA [8, 13]. This nuclease cleaves DNA targets that contain (i) a 20-nt complementary sequence with the crRNA guide, known as the protospacer, and (ii) an NGG sequence motif located immediately downstream of the protospacer, known as the protospacer-adjacent motif (PAM) [14, 15]. While these targeting rules are required to ensure that the response is specific against the virus, they also represent a liability to the host since phages carrying mutations in either the “seed” sequence of the protospacer (a critical region of 6-8 nt at the 3’ end) or PAM can escape the type II CRISPR defenses [16]. Our lab studies spacer acquisition in *Staphylococcus aureus* liquid cultures expressing the type II-A CRISPR-Cas locus of *Streptococcus pyogenes* SF370, after infection with the ϕNM4γ4 phage [17]. In this system, approximately 1 every 10^7^ non-immune infected cells survive through the incorporation of a new spacer, which occurs immediately upon infection and preferentially from the injected double-stranded DNA end of the phage [18]. The acquisition rate can be substantially increased in immune cells, where Cas9 cleavage of the viral DNA generates additional double-stranded DNA ends for the spacer acquisition machinery, a phenomenon known as “priming” [19]. While the viral population contains escaper phages that overcome the immunity mediated by a given guide RNA, these can lyse only bacteria harboring the cognate spacer, and phage propagation comes to a halt when they infect a host that contains a different spacer sequence [20]. Most staphylococci are immunized only once [17, 21], therefore the spacer diversity required for the neutralization of escaper phage is achieved at the population, as opposed to the cellular, level. A small fraction of staphylococci that acquire multiple spacers can also be found, however whether this feature is important for survival is poorly understood.

We hypothesized that spacer diversity within individual host cells should be important in communities inhabiting solid media (colonies). In this environment, bacteria that acquire new spacers grow into separated colonies that cannot mix to neutralize escapers. Therefore it would be reasonable to expect that the rise of escapers within colonies would quickly end with the annihilation of the host. This, however, is not the case, as many CRISPR-resistant colonies can be formed after infection of staphylococci with ϕNM4γ4 phage in solid media [17, 22]. Here we studied the formation of CRISPR-resistant colonies in top agar media. We found that the fate of the colony depends on the strength of the first spacer acquired. If the founder cell contains a spacer with a very low proportion of escapers in the phage population, the colony grows smoothly and maintains a stable coexistence with low levels of escaper phage. In contrast, when the viral population contains a high frequency of target mutations that avoid the type II CRISPR-Cas immunity mediated by the founder spacer, sectored colonies are formed, with each sector being composed of cells harboring additional spacers acquired after the founder spacer. In these colonies the new spacers are acquired through Cas9-mediated priming and are selected by the escaper phage, which lyses bacteria containing only the founder spacer and thus promotes the formation of sectored colonies. While the colonies originated by weak founder spacers struggle to rise when compared to those founded by strong spacers, they undergo a co-evolution with the phage that leads to the generation of spacer diversity and higher viral resistance. Our study reveals the versatility of type II CRISPR-Cas systems, which can respond with both single or multiple spacer acquisition modes to generate immunological diversity at the population or cellular level, respectively, and ensure survival in different environments.

## Results

### Bacteriophage-resistant colonies display two modes of spacer acquisition

After infection of a liquid culture of *Staphylococcus aureus* RN4220 cells [23] carrying the type II-A CRISPR-Cas locus of *Streptococcus pyogenes* SF370 [24] in the plasmid pC194 [25] with the ϕNM4γ4 phage [26], bacterial survival is achieved through the acquisition of spacer sequences from the viral genome [17]. Amplification of the CRISPR locus from the entire culture shows two PCR bands, one corresponding to bacteria that did not acquire new spacers and most likely succumbed to phage infection, and another corresponding to the acquisition of a single spacer (Fig. 1A). When we plated the infected culture to analyze the CRISPR locus of survivors, we confirmed that most cells acquired a single spacer, with occasional cases of two and three spacer acquisition events. Previous studies that performed next-generation sequencing of the expanded CRISPR array showed that thousands of different spacer sequences are acquired in this experimental system [17, 21]. This high diversity of CRISPR targets is required to neutralize the rise of “escaper” phages; i.e. phages containing mutations that avoid targeting by a given spacer sequence [20].

**Figure 1.**
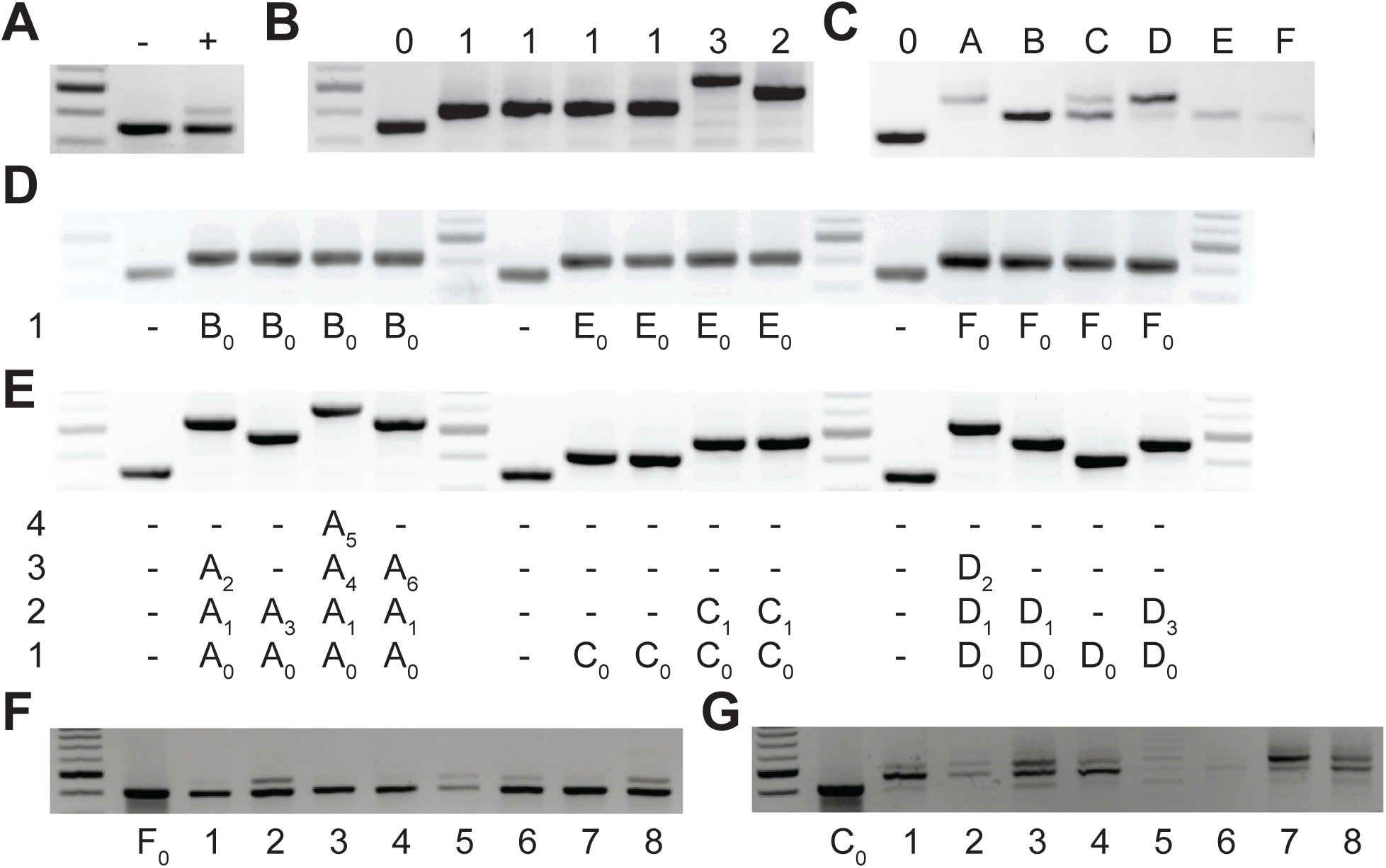
Bacteriophage-resistant colonies display two modes of spacer acquisition, determined by the sequence of the founder spacer. Agarose gel electrophoresis of the products of the amplification of the pCRISPR array present in staphylococci infected with ϕNM4γ4, using plasmid DNA templates extracted from: (**A**) liquid cultures with (+) or without (-) addition of phage; (**B**) colonies formed after plating the infected liquid cultures, with the numbers of new spacers indicated above; (**C**) bacteriophage-resistant colonies (labeled with different upper case letters) after infection in semi-solid (top agar) media, “0” indicates a no-spacer control sample; (**D**) colonies resulting from the re-streak of mono-spacer colonies shown in (**C**), with the spacer sequence determined after sequencing of the PCR product and a number indicating the position of the sequenced spacer(s) in the pCRISPR array; (**E**) same as (**D**) but after re-streak of multi-spacer colonies; (**F**) colonies resulting from infection in semi-solid media of cells containing pCRISPR with the founder spacer F (F_0_, pre-infection control); (**G**) same as (**F**) but for founder spacer C.

A key aspect of this mechanism of neutralization is the continuous mixing of the liquid culture, which ensures the infection of cells carrying different spacers that cannot be overcome by a particular escaper virus. We wondered whether and how escapers are neutralized in a more structured environment where phage and CRISPR-adapted cells cannot mix. To test this, we performed the same experiment described above, but in semi-solid agar plates, mixing bacteria and phage at a multiplicity of infection, MOI, of 2: 2×10^9^ staphylococci and 4×10^9^ ϕNM4γ4 particles. About 2,000 colonies grew after 48 hours of incubation, which we checked for spacer acquisition via PCR to find that some acquired a single spacer and others displayed a heterogenous composition (Fig. 1C); i.e. they contained cells with 1, 2, 3 and 4 spacers. To analyze the spacer content of different cells within these colonies, we re-streaked three of them and amplified and sequenced the CRISPR locus of four of the resulting colonies. We found that the original mono-spacer colonies were composed of staphylococci harboring the same spacer sequence (Fig. 1D and Supplementary sequences file). In contrast, the multi-spacer colonies contained cells with 1, 2 or 3 spacers (Fig. 1E and Supplementary sequences file). In addition, in all cases the sequence of the first acquired spacer was the same, indicating that the founder cell of the colony acquired a single spacer that allowed phage resistance, but as this cell divided more, different, spacers were added. To corroborate that the different cells of the colony originated from the same single-spacer ancestor, we employed a version of the type II-A CRISPR plasmid containing a randomized sequence upstream of the first repeat, that functions as a unique barcode for each plasmid [21]. We performed the same experiment using this plasmid and found a complete correlation between the barcode and first spacer sequence in each cell of the surviving colonies. This result shows (i) that the different spacers are not a consequence of the presence of different pCRISPR plasmids within the cells of the colony and (ii) that multi-spacer colonies are generated by a single cell that acquired a (founder) spacer (Fig. S1 and Supplementary sequences file).

### The sequence of the founder spacer determines the colony type

Next, we investigated the effect of the founder spacer on the expansion of the CRISPR array. We used plasmids isolated from one of the mono-spacer colonies or from one of the colonies that contained only the first acquired spacer obtained after re-streaking multi-spacer colonies. To eliminate any possible chromosomal mutations that could have played a role in the selection of the phage-resistant colonies, we re-introduced both plasmids into *Staphylococcus aureus* RN4220 cells and performed a modified version of the soft-agar infection experiment. Since these staphylococci contain a spacer that makes them immune to phage lysis, in order to mimic the conditions that lead to survival through spacer acquisition in the original experiment, we mixed only 2,000 CRISPR-immune cells (the previously determined number of CRISPR-survivors) with 1.3×10^9^ susceptible (without the CRISPR plasmid) staphylococci and 2.6×10^9^ ϕNM4γ4 particles (MOI = 2). As expected, after 48 hours of incubation, we obtained about 2,000 phage-resistant colonies on each plate and analyzed the spacer content of 8 of them. We found that the colonies obtained from the mono-spacer founder plasmid did not acquire additional spacers (Fig. 1F). In contrast, the surviving colonies produced from the multi-spacer founder plasmid harbored many additional spacers (Fig. 1G). This result indicates that the first spacer acquired by a CRISPR-adapted cell determines the spacer content of its progeny during the formation of the colony.

### CRISPR expansion involves priming by the first acquired spacer

To analyze in more detail the spacer content of the surviving population, we isolated 26 individual colonies from a replica of the experiment described in Figure 1C, amplified their CRISPR array (Fig. 2A) and subjected the PCR products to next-generation sequencing (NGS). We found 10 colonies that produced >98 % of single-spacer reads (Fig. 2B and Supplementary sequences file) and therefore they belong to the mono-spacer category, also evident from the results of Figure 2A. The remaining 16 colonies displayed a variable fraction of multi-spacer reads, ranging from 5 to 85 % (Supplementary Data File 1). We used these data to determine whether “priming” is involved in the sequential acquisition of new spacers. During primed CRISPR adaptation the frequency of spacer acquisition is significantly increased by the presence of a pre-existing targeting spacer [27]. When we use our experimental system to investigate the infection of cells in liquid culture, either partial or complete Cas9 cleavage of the phage genome directed by a failing or efficient, respectively, first acquired spacer results in subsequent acquisition of additional spacers, most of them from regions flanking the Cas9 target site [19]. To test for this feature, we extracted 1,711 spacer pairs from the NGS data and measured the first-to-second spacer distance (Fig. 2C). We found that, similarly to our previous results in liquid cultures, there is a marked enrichment of second spacers derived from the 500 bp region adjacent to the target of the first spacer. This result suggests that primed, or cleavage-mediated, spacer acquisition plays an important role to create multi-spacer cells and colonies.

**Figure 2.**
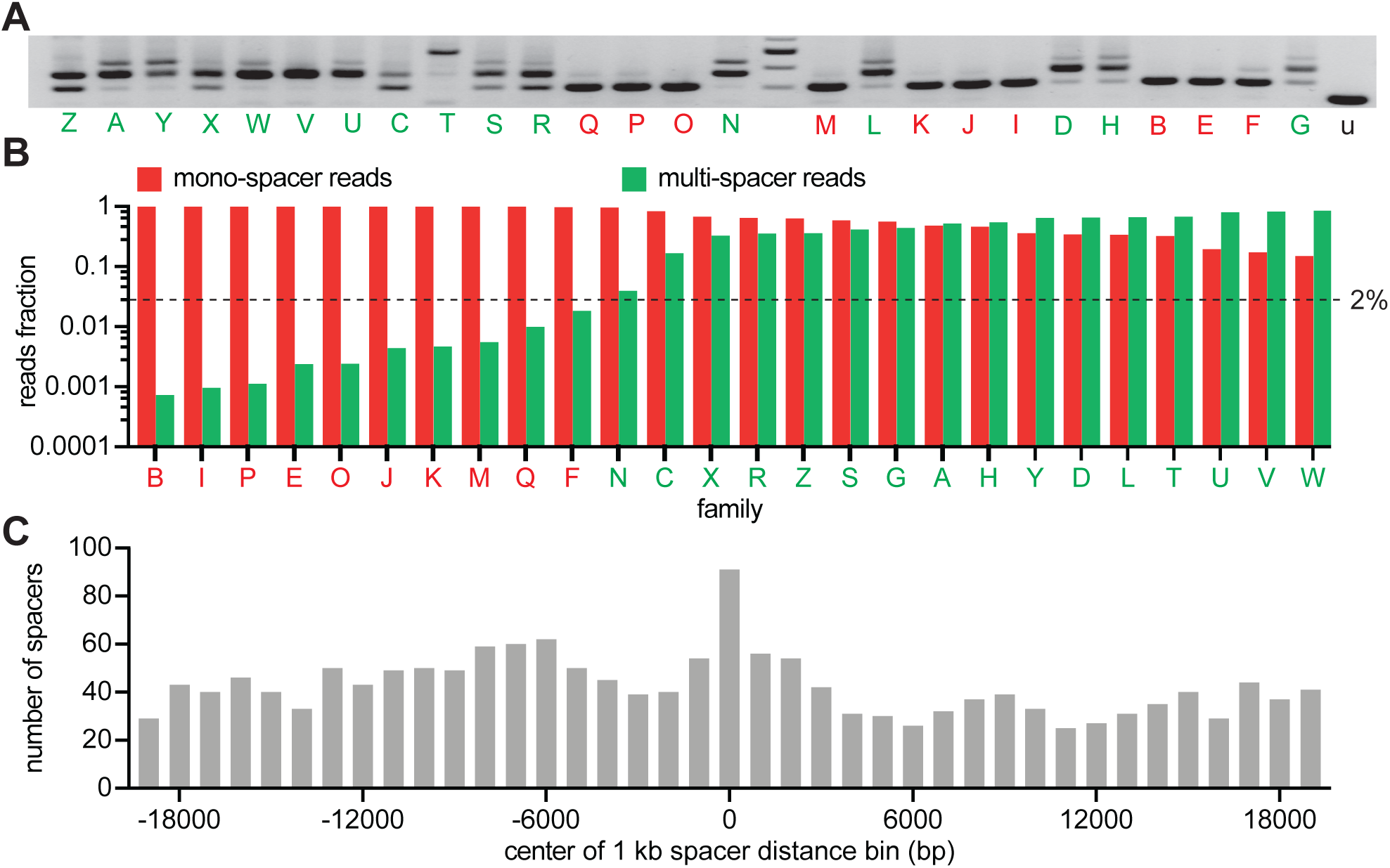
CRISPR expansion involves priming by the first acquired spacer. (**A**) Agarose gel electrophoresis of the products of the amplification of the pCRISPR array present in staphylococci infected in semi-solid media with ϕNM4γ4, using plasmid DNA templates extracted from 26 different surviving colonies; red and green letters: mono- and multi-spacer colony names. (**B**) Fraction of the reads obtained after NGS of the PCR products shown in (**A**) containing either a single or multiple spacers (red and green bars, respectively). Dashed line indicates the 2 % value, the minimal fraction of multi-spacer reads that leads to multiple PCR products after amplification of the pCRISPR array. (**C**) Distance between the targets in the ϕNM4γ4 genome specified by the first and second spacers acquired after infection of staphylococci carrying pCRISPR; obtained from analysis of NGS data. The number of different second spacers within 1-kb bins of the ϕNM4γ4 genome are shown; the position of first spacer acquired in each array is set as 0 kb.

### The ability of phage to escape targeting by the founding spacer determines colony heterogeneity

Interestingly, the above results showed that mono-spacer colonies also contained a low proportion of cells harboring multiple spacers, only detected by next-generation sequencing but not by PCR of the CRISPR array. This finding demonstrates that multiple spacer acquisition occurs in both colony types, most likely through priming as shown above. Thus, the generation of multi-spacer colonies can be explained by at least two non-exclusive mechanisms. One possibility is that the founder spacer mediates very efficient priming that leads to the generation of high numbers of multi-spacer cells as the colony forms. Another scenario is that additional spacer acquisition through priming mediated by the founder spacer is equally infrequent, but the viral population contains a high frequency of escaper phages that can evade the immunity mediated by the founder spacer. This results in the positive selection of the few cells that contain an extended CRISPR array, leading to the formation of multi-spacer colonies. In this scenario, founder bacteria that acquired spacers with a low escape rate in the viral population grow largely unchallenged; cells harboring additional spacers are not selected and a mono-spacer colony is formed. We tested this hypothesis with three experiments. First, we measured the escape rate of different founder spacers by calculating the fraction of escapers within the phage population. We found that 8/9 founder spacers from multi-spacer colonies showed high rates of escape, where 1 in 10^6^-10^7^ phages were able to evade type II CRISPR-Cas immunity (Fig. 3A). In contrast, founder spacers (10 tested) of mono-spacer colonies displayed a much lower rate of escape, ranging from undetectable to a maximum of 10^−8^. Sequencing of the different targets confirmed that the escaper phages contained seed or PAM mutations (Fig. S2). Second, we looked for the presence of escaper phage in the colonies that resulted from infection of mono-spacer founder cells in the experiment shown in Figure 1F and were unable to detect phages that could bypass spacer F_0_ immunity (Fig. S3A). Third, we measured the presence of phages that specifically evade targeting by the founder spacer within the surviving colony. To do this, we resuspended two multi-spacer phage-resistant colonies in media and centrifuged them to separate cells from soluble phage particles. The pelleted staphylococci were re-streaked and checked for the number of acquired spacers to isolate cells with only one, two or three spacers. Lawns of these bacteria were infected with 10-fold dilutions of phage from either the wild-type stock or isolated from each of the colonies (Fig. 3B and Supplementary sequences file). We found that staphylococci harboring only the founder spacer were readily infected by the phage isolated from their own colony, but not by the second phage. The acquisition of an additional spacer drastically reduced the propagation of the phage, and the insertion of the third spacer drove plaque formation below the limit of detection of the assay.

**Figure 3.**
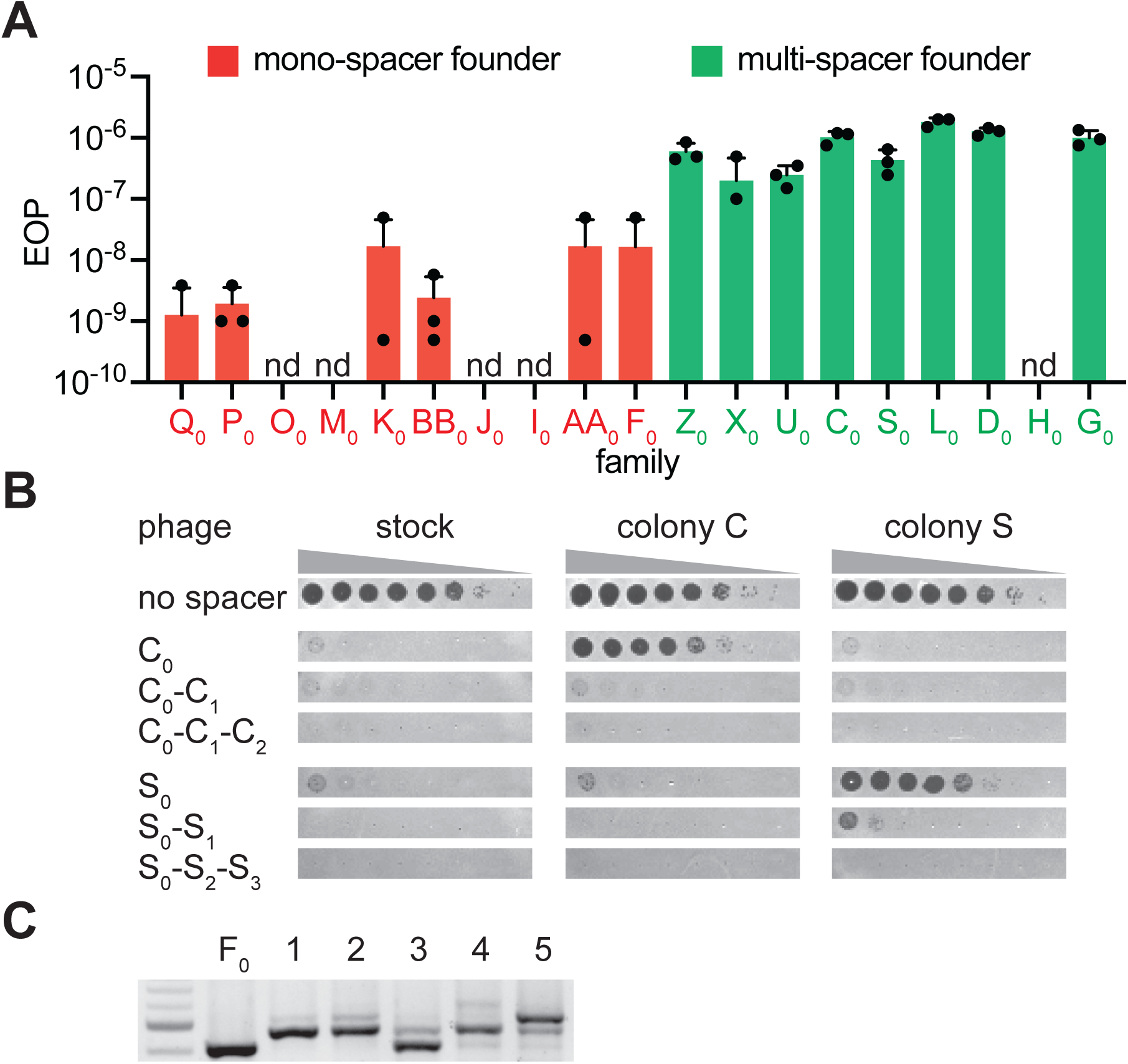
The ability of phage to escape targeting by the founding spacer determines colony heterogeneity. (**A**) Efficiency of plaquing (EOP), calculated as the number of ϕNM4γ4 plaques on the test strain relative to the total number of phage particles in the stock. Different mono-spacer (red) and multi-spacer (green, carrying only the founder spacer without additional ones) strains were tested. Mean ± STD of 3 biological replicates (black dots) are reported. (**B**) Detection of plaques present in 10-fold serial dilutions of either ϕNM4γ4 phage stock or phage isolated from multi-spacer colonies founded by spacers C or S, spotted on lawns of non-CRISPR staphylococci (“no spacer”) or carrying pCRISPR plasmids with an increasing spacer content. (**C**) Agarose gel electrophoresis of the products of the amplification of the pCRISPR(spacer F) array present in five colonies that survived infection in semi-solid media with ϕNM4γ4 and spacer F escaper phage.

Next, we tested if the high proportion of escapers of a founder spacer is the cause for the formation of multi-spacer colonies. If this is the case, founder spacers from mono-spacer colonies would also form multi-spacer colonies when escaper phage are artificially added to the phage stock. We isolated a phage that can overcome targeting by spacer F_0_ (Fig. S2), a mono-spacer colony founder, and repeated the experiment of Figure 1F infecting with a mixture of wild-type and escaper phage. We previously measured that the frequency of escape of spacer F_0_ is ∼ 10-8 (Fig. 3A); we added escaper phage to raise this rate to 10^−5^. To eliminate the possibility of escaper-mediated primed spacer acquisition [19] (different from the priming elicited by cleavage of wild-type phage that we described above) that could confuse the interpretation of the results, we used an escaper harboring mutations in both the seed and PAM target sequences (Fig. S2), which should not be recognized by Cas9 loaded with the F_0_ crRNA to elicit priiming. As opposed to the previous results, the increase in the proportion of escaper phage generated multi-spacer colonies from this otherwise mono-spacer colony founder (Fig. 3C) and contained phage that escaped the immunity provided by the F_0_ spacer (Fig. S3B). Together, these results demonstrate that the escape rate of the founder spacer determines the spacer content of the colony, with high levels of escape leading to the formation of multi-spacer colonies.

### Multi-spacer colonies have impaired growth

Given that primed spacer acquisition is a relatively rare event [19], the majority of cells in a developing colony contain only the founder spacer. Therefore, a high rate of escape of this spacer would lead to the predation of most cells in the bacterial community by target mutant phages, possibly even abrogating the formation of the colony. To corroborate this, we quantified the number of colonies obtained in the experiments shown in Figures 1F, 1G and 3C. Indeed, whereas about 1,000 of the 2,000 staphylococci harboring the mono-spacer founder F_0_ added to the top agar survived infection, only about 500 cells containing the multi-spacer founder C0 were able to form colonies (Fig. 4A). To investigate the role of spacer acquisition in the formation of the colonies, we also infected cells containing pCRISPR(Δ*cas2*), which are unable to acquire new spacers [17]. As expected, this mutation profoundly affected the growth of the C_0_-founded colonies, which rely in the addition of new spacers to survive escaper phages. In contrast, the Δ*cas2* genetic background did not have a big impact in the rise of F0-founded colonies (Fig. 4A). However, when the F_0_ cells where infected in the conditions described for the experiment in Figure 3C, i.e. with a phage population carrying a high proportion of escapers that evade the immunity provided by this spacer, colony formation mimicked the results obtained for the C_0_ founder cells infected with a regular phage stock (Fig. 4A). These observations confirm our previous results and highlight the role of escaper phage in the formation of different types of CRISPR immune colonies.

**Figure 4.**
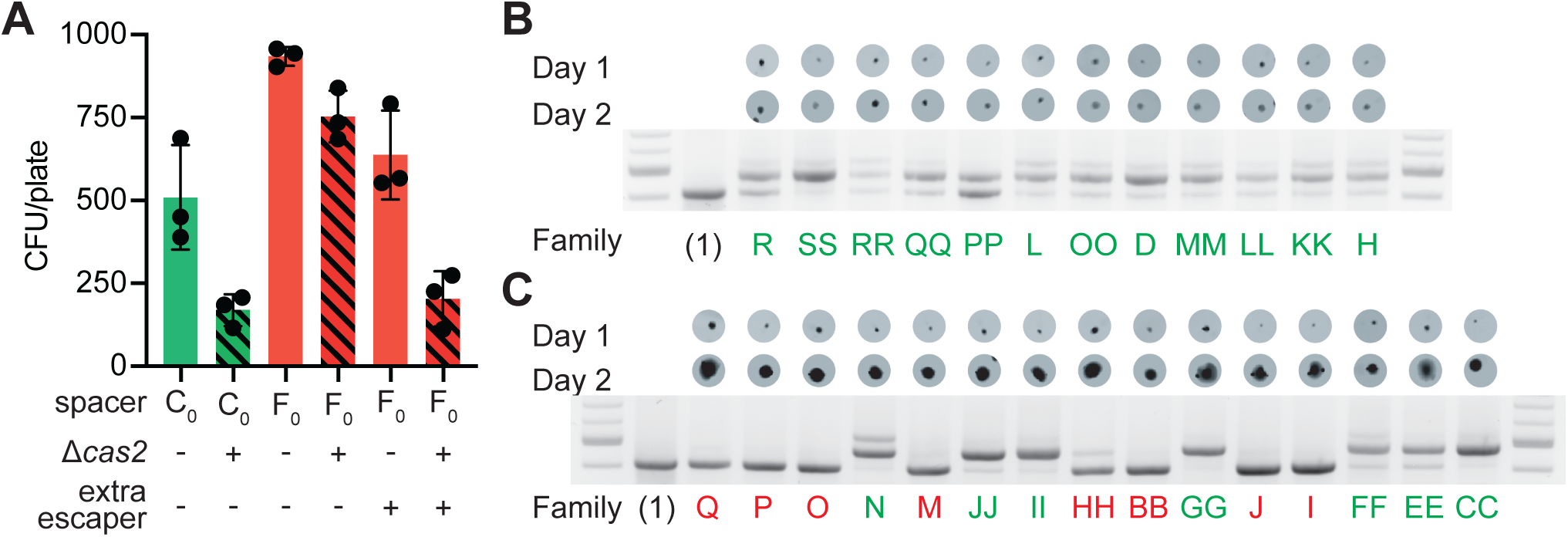
Multi-spacer colonies have impaired growth. (**A**) Enumeration of colony forming units (CFU) obtained after infection of staphylococci carrying pCRISPR or pCRISPR(Δ*cas2*) plasmids (clear or dashed pattern bars), containing spacer C or F (red or green bars, respectively), with ϕNM4γ4 or ϕNM4γ4 also containing spacer F escaper phage. (**B**) Images of bacteriophage-resistant colonies 1 or 2 days after infection with ϕNM4γ4 phage, and PCR analysis of the spacer content in their pCRISPR plasmid. (1), one spacer control. (**C**) Same as (**B**) but with colonies that experienced growth over time; their letter name colored according to the spacer content: red, mono-spacer; green, multi-spacer.

While many of the staphylococci that acquire a weak spacer cannot develop a colony, for those that do form a colony it is expected that escaper phages would decimate most of the cells that harbor the founder spacer and only the multi-spacer cells should be able to grow. If this is true, multi-spacer colonies should display a slow growth rate. To test this, we performed the experiment of Figure 1C and imaged plates at 24 hours and 48 hours after phage infection. We followed 27 colonies displaying similar sizes at 24 hours, and found that 12 did not change in size (Fig. 4B) and 15 that were larger (Fig. 4C) at 48 hours. Each colony was then checked for the expansion of the CRISPR locus. As hypothesized, all colonies that were unable to grow (12/12) acquired multiple spacers (Fig. 4B and Supplementary sequences file). On the other hand, colonies that increased in size over time contained both a single (8/15) or multiple (7/15) spacers (Fig. 4C and Supplementary sequences file).

### Multi-spacer colonies display sectored morphologies

We decided to examine in more detail colonies the mono- and multi-spacer colonies that displayed detectable growth (Fig. 4C). When such colonies were photographed at higher magnification, we found two types of morphologies: smooth and sectored (Fig. S4A). Analysis of the spacer content of several of these colonies revealed that while the smooth colonies contained a single spacer (Fig. 5A and S4B-D and Supplementary sequences file), the sectored ones harbored multiple spacers (Fig. 5B and S4E-G and Supplementary sequences file). Moreover, analysis of isolated sectors showed that they contained different multi-spacer families with the same founder spacer (Fig. 5B and S4G and Supplementary sequences file). To investigate the role of escaper phage in the formation of sectored colonies, we performed the experiment of Figure 3C, infecting mono-spacer founder F cells, both capable (wild-type) or unable to acquire new spacers (Δ*cas2*), with an excess of escaper phage. Indeed, exposure to high concentrations of escaper phage, but not to the wild-type phage stock alone, resulted in the formation of sectored colonies (Figs. 5C and S5A). Moreover, in the absence of spacer acquisition the addition of escaper phage resulted in the formation of translucent Δ*cas2* colonies, most likely formed before the rise of escapers and then disintegrated. To corroborate these results, we grew mono-spacer, smooth, colonies in the absence of phage for 24 hours and then added escaper phage on top of them (Fig. 5D). We also added PBS as a negative control or lysostaphin, a peptidoglycan hydrolase [28] (Fig. S5B), as control for colony lysis. After 24 hours of incubation we observed that PBS did not alter the morphology of the colonies; i.e. they remained smooth. As expected, lysostaphin lysed the cells and produced translucent colonies. In contrast, the addition of exogenous escaper phage altered the morphology of the colonies, from smooth to sectored. These results demonstrate that founder spacers determine the morphology of the colony they originate, most likely by affecting the co-evolution dynamics that follows the rise of CRISPR-adapted hosts.

**Figure 5.**
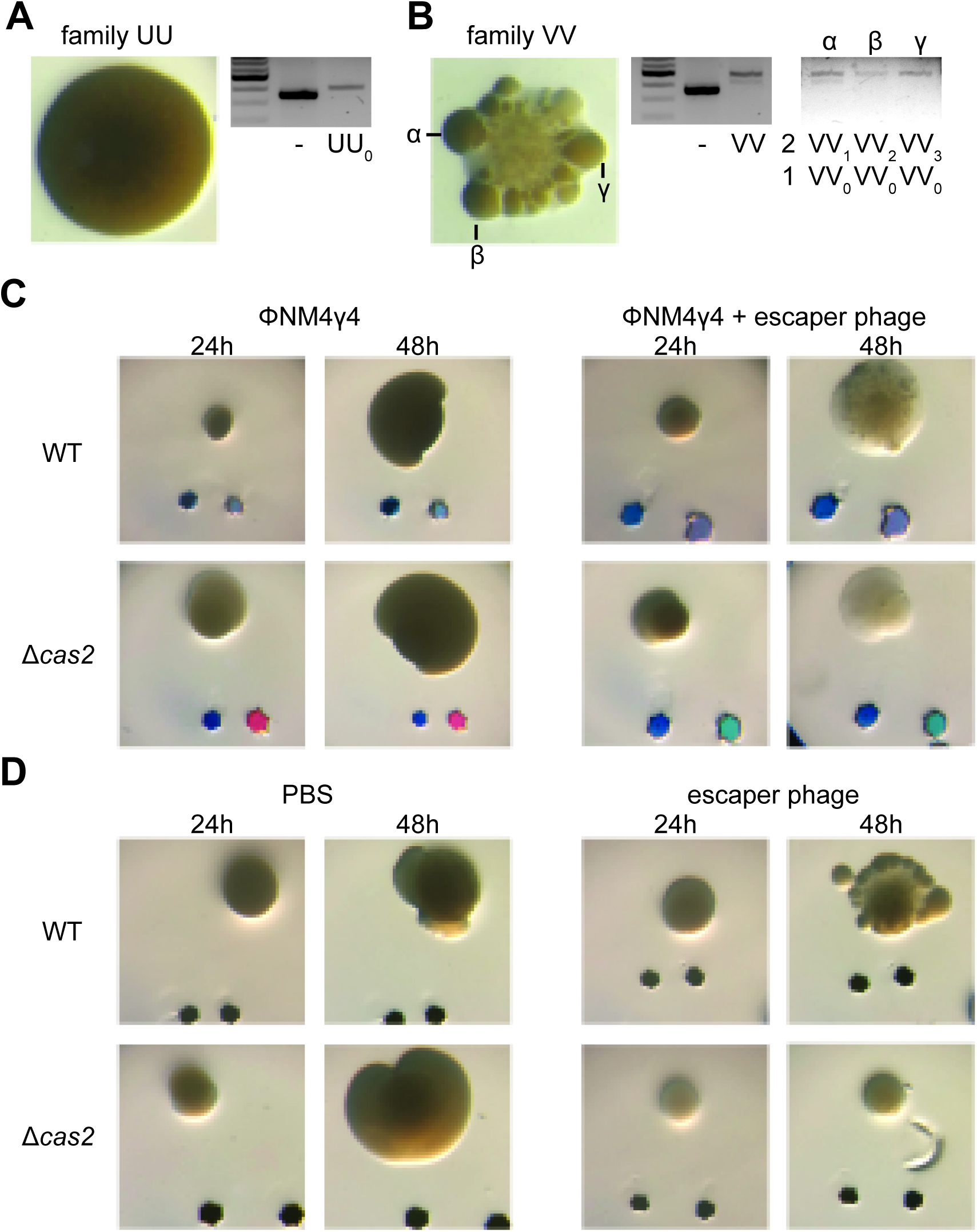
Multi-spacer colonies display sectored morphologies. (**A**) Image of a smooth colony (containing the founder spacer UU) as well as the gel agarose analysis of PCR products obtained after amplification of its pCRISPR array; (-) shows amplification of pCRISPR, a no-spacer control. (**B**) Same as (**A**) but for the sectored colony founded by spacer VV; and also showing the gel agarose analysis of PCR products obtained after amplification of its pCRISPR array present in three different sectors (α,β,γ) where 1 and 2 refer to the first and second position, respectively, of the sequenced spacer(s) in the pCRISPR array. (**C**) Images of representative colonies grown 24 or 48 hours after top agar infection of wild-type or Δ*cas2* mono-spacer founder F cells, with ϕNM4γ4 containing or not additional spacer F escaper phage. Glitter markers are shown to normalize both the position as well as the size of the image at different times. (**D**) Images of representative colonies of wild-type or Δ*cas2* mono-spacer founder F cells grown for 24 hours in the absence of phage, when a drop of either PBS or spacer F escaper phage was added on top and a second image was obtained 24 hours after. Glitter markers are shown to normalize both the position as well as the size of the image at different times.

### Host-phage co-evolution within multi-spacer colonies leads to increased resistance

The model described above suggests that cells within the multi-spacer colonies, originally founded by a weak spacer, co-evolve over time with the phage to limit its ability to infect. Strong founder spacers, in contrast, should quickly neutralize the phage. To test these hypotheses, we performed a time-shift experiment that evaluates the resistance of bacterial samples over time against phage from past, concurrent, and future time points [29, 30]. We repeated the experimental setup used in Figures 1F (mono-spacer founder F) and 1G (multi-spacer founder C), collecting 12 surviving colonies at 24h, 36h, and 48h post-infection and isolating and amplifying phage from each of them Finally, each colony and their founders were grown in liquid cultures and used to seed top agar media, on which 2 μl of stock phage (which has not co-evolved with CRISPR-resistant staphylococci and was used to obtain the “time zero” datapoint) or phage isolated from each colony was spotted. After overnight incubation at 37 °C, we evaluated the resistance of the bacteria on the top agar as follows: a phage spot that did not generated an growth inhibition zone was taken as ‘full resistance’ and given a score of 1; complete inhibition of growth was determined to be ‘no resistance’ and given a score of 0. Partial inhibition was given a score of 0.5. The resistance scores for the bacteria against these 12 phage populations of a given time-point were averaged for each host, with the exception of the time zero datapoints, obtained with only the stock phage (Supplementary data file).

To evaluate whether colonies of a given time-point xgained or lost resistance over time, we calculated the average resistance of a colony against all the phage from the same time point and plotted the score averages for each colony over time (Fig. 6 and Supplementary data file). We found that the mono-spacer founder F is initially resistant to the stock phage, but experiences a small decrease in immunity (from a score of 1 to 0.83) against all “future” phages (Fig. 6A), demonstrating that the co-evolution of the phage with cells carrying this spacer generates a low number of escapers that increases over time. This co-evolution also increases the resistance of post-infection bacteria collected at 24, 36 and 48 hours (Figs. 6B-C), leading to a stable co-existence of phage and resistant bacteria at the end of the experiment. In contrast, the results obtained after following the infection of spacer C hosts revealed a more drastic co-evolution dynamics. First, founder cells showed a marked susceptibility to “future” phages that co-evolved over time (from a score of 1 to 0.5, Fig. 6E), an observation in agreement with previous results showing that spacer C is more likely to encounter escaper phages (Fig. 3A). Second, co-evolution increases the overall resistance of colonies collected at 24 hours after infection (Fig. 6F) and then leads to complete resistance at 36 and 48 hours post-infection (Figs. 6G-H). Therefore, the increased pressure that founder spacer C faces from escaper phages seems to both accelerate and strengthen the evolution of resistance.

**Figure 6.**
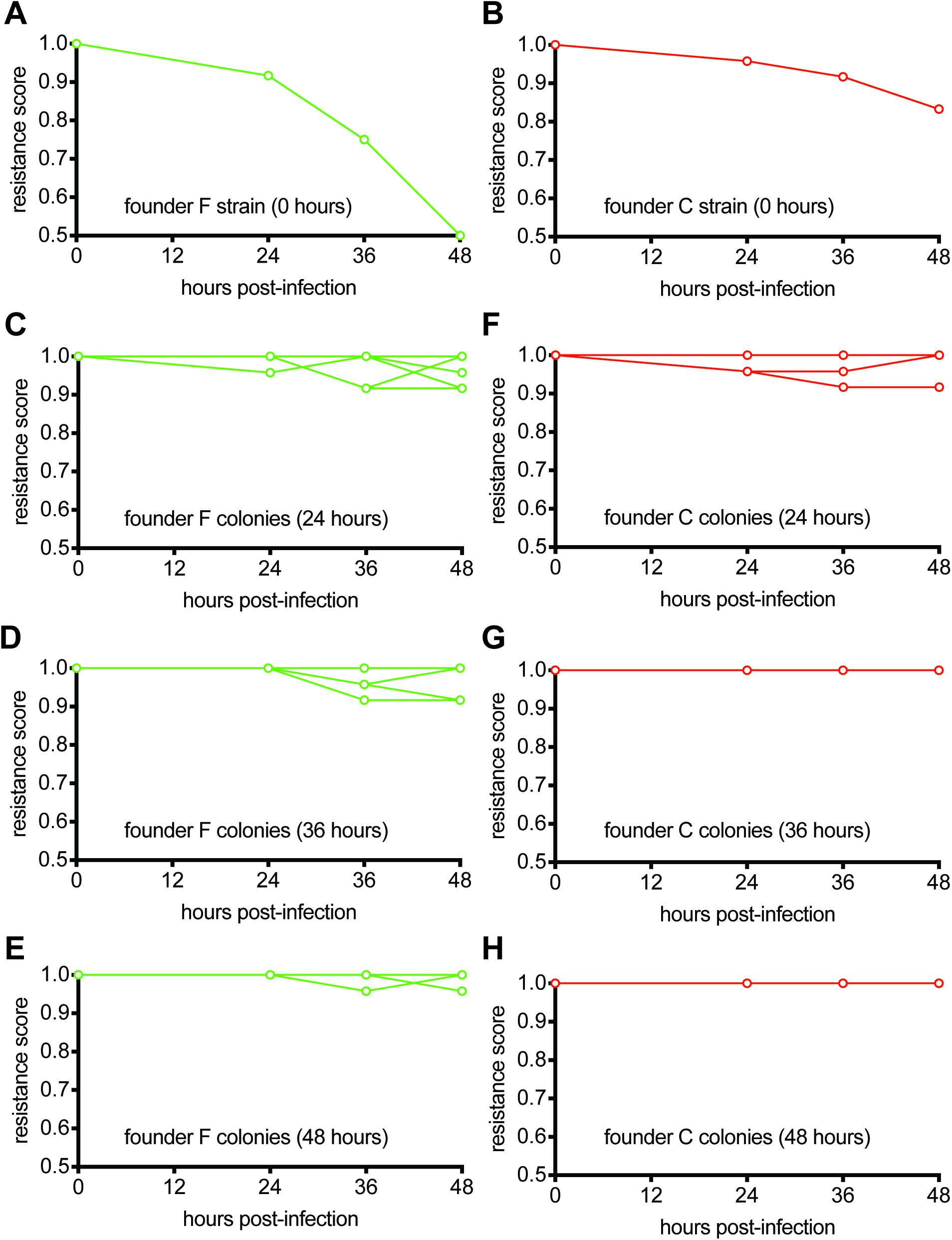
Host-phage co-evolution within multi-spacer colonies leads to increased resistance. (**A**) Resistance score of staphylococci carrying the pCRISPR programmed with founder spacer F to the stock ϕNM4γ4 phage (time 0) or the phages isolated from colonies that survived 24, 36 and 48 hours after infection. (**B**) Same as (**A**) but for cells with pCRISPR programmed with founder spacer C. (**C**-**E**) Same as (**A**) but showing the resistance scores for 12 different colonies isolated at 24 (**B**), 36 (**C**) and 48 (**E**) after infection. to the stock ϕNM4γ4 phage (time 0) or the phages isolated from colonies that survived 24, 36 and 48 hours after infection. (**F**-**H**) Same as (**C**-**E**) but for cells with pCRISPR programmed with founder spacer C.

## Discussion

Here we studied the formation of resistant colonies during the type II CRISPR-Cas immune response. CRISPR immunity starts with the acquisition of a spacer sequence from the invading phage, which confers resistance to the cell. When infection occurs in solid media, each of these resistant bacteria can form a colony. We found that the strength of the immunity provided by the spacer acquired in the founder cell determines the nature of the virus-host co-evolution that occurs during the formation of the colony. This co-evolution in turn defines the spacer diversity, growth rate, morphology and overall resistance of the colony. Founder cells harboring a weak spacer, which immunity can be overcome by a relatively high number of target mutant phages in the viral population, engage in a dynamic co-evolution with the virus. These cells suffer the predation of escaper phages and struggle to grow; members of the colony in formation that carry additional spacers are selected and their growth leads to the formation of micro-colonies, cellular conglomerates with the increased spacer diversity necessary to limit the propagation of escaper phages. On the other hand, the phage is largely contained by founder cells harboring strong spacers, and little viral-host co-evolution occurs. Escapers do not rise, cell growth is not limited, and smooth, mono-spacer colonies are formed. Interestingly, the initial and final levels of resistance of the colonies are reverted. The virus-host co-evolution mediated by the CRISPR-Cas immune response enables the conversion of a genotype that initially confers a high level of susceptibility to the host (a weak founder spacer) into a new genotype that not only promotes survival from co-evolving phages, but also creates a completely resistant host.

In theory, there are two known priming mechanism that can explain how additional spacers are acquired within the colonies. Both depend on the phage that is attacked by Cas9 (spacers are acquired from the DSBs generated by the nuclease [19]) but differ in the nature of the target sequence. In one case, wild-type phage is cleaved within already immune cells, which acquire extra spacers in anticipation of the rise of escapers. In the other case, a low proportion of escape phages containing target mutations that allow low levels of Cas9 cleavage, not sufficient to confer immunity, lead to additional spacer acquisition within susceptible hosts. Although both situations are formally possible and not mutually exclusive, the last scenario is highly unlikely. This is because the low level of cleavage makes the host susceptible to predation by the escaper phage and therefore the fraction of cells that survive infection and acquire additional spacers is very low, about 1 cell per 10^7^ to 10^6^ in our experimental system [19] (this fraction could be even lower in our experiments with artificially increased levels of spacer F escaper phage, which carries mutations in both the seed and PAM sequences and therefore is even more refractory to occasional Cas9 cleavage). Given that a developing colony contains an even lower number of cells and that we commonly observed the acquisition of more than two spacers (Fig. 1E), it would be mathematically impossible for CRISPR adaptation to occur not only once but twice or three times within the same host. Therefore, we favor the first scenario, in which a fraction of the immune cells exploit Cas9 target cleavage not only to stop infection but also for the acquisition of several new spacers. In contrast to the occasional cleavage of escaper phages, this mode of spacer acquisition can lead to the addition of several spacers within a short time, since the incorporation of a second spacer mediates two Cas9 cleavage events and therefore duplicate the chances of acquisition of a third spacer. We think that this occurs in a small fraction of the cells within all developing colonies, which are further amplified or not depending on the number of escaper phage that evade the founder spacer in the colony.

Interestingly, the most detailed analysis performed in this study, i.e. the detection of expanded CRISPR arrays via next generation sequencing, showed that the of fraction of multi-spacer loci within each colony was highly variable (Fig. 2B). There are two non-mutually exclusive factors that could affect this value: the escape frequency and/or the priming efficiency of the founder spacer. We believe that while the first factor impacts more profoundly the fraction of multi-spacer cells in colonies that, when analyzed with less powerful methods such as colony PCR (Fig. 2A) and microscopy (Figs. 5A-B), display a multi-spacer and sectored phenotype, the second factor is determinant for the different proportions of multi-spacer cells in colonies that appear as mono-spacer and smooth. If high efficiency of priming was responsible for the formation of colonies with high proportion of multi-spacer cells, it would lead to the immediate acquisition of additional spacers by a high proportion of the infected cells in the developing colony. Such mechanism, however, should result in the formation of smooth multi-spacer colonies, as it would achieve an early neutralization of the founder spacer escapers and allow the unchallenged growth of the colony. Therefore, we believe that the wide distribution of the fraction of multi-spacer cells within sectored colonies reflects the different escape frequencies for each founder spacer. While we defined founder spacers as weak or strong, in truth the escape frequencies have many different values depending on the region of the phage genome targeted by each spacer. At one end of the spectrum will be the spacers that target viral protospacers for which any modification of the seed or PAM sequence would lead to complete loss of phage viability. At the other end will be the spacers that match a protospacer that when their seed or PAM sequences are changed, the mutation enhances viral propagation. Many escaper mutations, however, will fall in between these two scenarios, and will include mutations that only decrease or are neutral for the fitness of the phage. This will produce a wide range of escaper mutation frequencies in the viral population that will lead to different levels of selection and accumulation of multi-spacer cells within sectored colonies.

On the other hand, while founder spacers of smooth colonies also display different escape frequencies (Fig. 3A), these are not high enough to affect the growth of the cells within the colony (Fig. 4C). Once cells reach stationary phase, they can no longer be infected by the escaper phage (staphylococci in this growth stage are refractory to infection by ϕNM4γ4 and related bacteriophages) and therefore they cannot propagate and promote the expansion the small fraction of multi-spacer cells present in smooth colonies. Instead, low levels of escaper phage remain in these colonies, even after 48 hours of the initial infection (Fig. 6E). We determined an empirical value of 2 % as the maximum fraction of multi-spacer array that can produce a mono-spacer and smooth phenotype (Figs. 2A-B). Since the multi-spacer members of these colonies are not important to limit the propagation of the escaper phages, their fraction is most likely a result of the different priming efficiencies of their founder spacers.

In our experimental system, infection of liquid cultures leads to a multi-spacer population composed of mono-spacer cells [17, 21]. In these studies, the initial spacer diversity neutralizes most escape variants present in the viral population and allows the fast growth of the culture. When it reaches stationary phase, it becomes immune to further infection and viral-host co-evolution stops. However, experiments with *Streptococcus thermophilus* DGCC7710 (a natural host of type II CRISPR-Cas systems) in which infected liquid cultures are artificially diluted before cells can reach stationary phase, also resulted in the accumulation of multi-spacer CRISPR loci in the population. [29, 31, 32]. It has been shown that under a dilution regiment, *S. thermophilus* and its phage 2972 both increase their resistance and infectivity, respectively, a result that demonstrates the existence of a co-evolutionary arms race [29]. After nine days of the initial infection phages lose their infectivity and the surviving cells contain multiple spacers. We believe that these results are in line with our findings, since dilution (i) prevents the culture from accumulating phage resistant, stationary phase cells, and (ii) reduces the spacer diversity of the culture and leads to the selection of multi-spacer that can contain the escapers. Therefore, even if our experimental system is not found in nature, there are reasons to believe that our findings could generally apply to the co-evolution in solid media of prokaryotes harboring type II CRISPR-Cas systems and their phages. Studies designed to follow the formation of CRISPR-resistant colonies using non-engineered hosts and phages, and, more importantly, the investigation of the evolutionary dynamics within natural ecosystems, remain to be performed.

In summary, we found that the acquisition of multiple spacers within a single CRISPR array compensates for the inefficient defense provided by an initial weak spacer, in conditions of structured growth where this compensation cannot be offered by a strong neighbor cell. Therefore the type II CRISPR-Cas immune response is able to provide robust immunity through the generation of either a diverse population composed of weak and strong members or a diverse cell harboring weak and strong spacers.

## Methods

### Bacterial strains and growth conditions

Cultivation of *S. aureus* RN4220 [23] or OS2 [33] was carried out in tryptic soy broth (TSB) at 37°C. *S. aureus* media was supplemented with chloramphenicol at 10 μg/ml to maintain pCRISPR plasmids.

### Spacer acquisition in liquid cultures

Overnight cultures of *S. aureus* RN4220 containing pCRISPR or pCRISPR(Δ*cas2*) (pWJ40 or pRH61, respectively [17]) were diluted 1:100 in 20 mL of BHI and grown for 4 hours shaking at 37°C. The optical density value at 600 nm (OD_600_) was measured and used to calculate the colony forming units (CFU) per μL. For the infection, 1.3 billion CFU of pCRISPR- or pCRISPR(Δ*cas2*)-containing cells were mixed with 2.6 billion plaque forming units (PFU) of ϕNM4γ4 bacteriophage [26] for a starting MOI of 2. This mixture was added to 65 mL of 50% Heart Infusion Broth (HIB) supplemented with 5 mM CaCl_2_ and incubated under shaking conditions at 37°C for 48 hours. Cells were stored by collecting 1 ml of the culture, which was mixed with 50% glycerol and stored at 4°C or frozen at -20°C. Individual colonies were isolated by spreading the culture on a TSA plate.

### Spacer acquisition in semi-solid media (top agar)

Infections in top agar were performed with almost the same conditions as those in liquid culture, except that the phage mixture was added to 5 mL of melted 50% Heart Infusion Agar (HIA top agar) supplemented with 5 mM CaCl_2_ instead of added to liquid media. The top agar was then poured onto a plate containing solidified Tryptic Soy Agar (TSA). Plates were then incubated at 37°C for 48 hours. Each colony was picked and resuspended in 30 μL of TSB and stored at 4°C or frozen at -80°C with 50% glycerol. To isolate individual cells from the colony, the adapted colony was diluted in TSB and streaked on TSA plates.

### Simulation of spacer acquisition in semi-solid media

To recreate the infection conditions encountered by the founder cell (an isolated CRISPR-immune cell suffering infection by a very high number of phages; the result of their exponential propagation within sensitive hosts that were not able to acquire spacers), we mixed 2,000 CFU of the founder cells (harboring pCRISPR with the addition of the founder spacer) with 1.3 billion CFU of pCRISPR(Δ*cas2*) cells before adding phage and mixing with top agar.

### CRISPR array amplification

To evaluate the size of the CRISPR array, we mixed 2 μl of the resuspended colony to 30 μL of colony lysis buffer with 50 ng/μl lysostaphin [17]. This mixture was boiled at 98°C for 10 minutes and cooled down at 37°C for 20 minutes to lyse the cells. To amplify the CRISPR array we used 1 μl of the supernatant as template for a TopTaq PCR amplification (Qiagen) with primers H54 and H237 (see Supplementary sequences file). The resultant PCR amplicons were then analyzed on 2% agarose gels. To further characterize individual sister cells within the colony, these PCR products were submitted for Sanger sequencing.

### Next-generation sequencing

pCRISPR plasmids were isolated from 10 μL of the frozen colonies with modified QIAprep Spin Miniprep Kit protocol as previously described [18]. We used 2 μL of the plasmid preparation as template for PCR amplification with Phusion DNA Polymerase (Thermo) using primers NP389 and NP390 (see Supplementary sequences file). Amplicons were given to the Rockefeller University Genomics Core for library preparation and Illumina Hi-Seq Sequencing. The data was analyzed using Python: spacer sequences were extracted ordered in according to their position within the array and aligned to the ϕNM4γ4 reference genome.

### Quantification of phages escapers for different founder spacers

Top agar lawns of each founder cell were made by mixing 100 μL of an overnight culture with 5 mL of HIA top agar supplemented with 5 mM CaCl_2_. We then plated 2 μL of a serially diluted phage stock on the lawn and incubated the plate at 37°C. After 24 hours we were able to count PFU and calculate the escaper rate for a particular founder spacer as the ratio to the total PFU count of the phage stock. DNA from individual escaper plaques was isolated according to previously described techniques [26]. The target region of each founder spacer was then amplified using the following oligonucleotides: Founder Z: H463, H493, Founder X: H40, NP255, Founder G: NP170, AV363, Founder S: NP331, H462, Founder D: NP180, H485, Founder Q: W1051, NP182, Founder P: NP279, W1087, Founder K: H501, H471, Founder BB: H122, H135, Founder F: AV462, H453, Founder AA: H477, NP313, Founder Y: NP265, H613, Founder C: H122, H135, Founder L: H501, H471; and the PCR products sent for Sanger sequencing.

### Isolation of phage from colonies

To propagate the phage and increase its titer, 10 μL of the supernatant from the resuspension of a resistant colony were mixed with 100 μL of an overnight culture of *S. aureus* OS2, an erythromycin-resistant non-immune strain, and 5 mL of HIA top agar supplemented with 5 mM CaCl_2_ instead. The mix was plated on erythromycin-containing TSA plates to ensure that host bacteria were killed. After 24 hours of incubation at 37°C phages were purified by filtration as previously described [17].

### Challenge of founder spacer F cells with additional escaper phage

First an escaper phage containing mutations in both the seed and PAM sequences of the spacer F target was isolated (Fig. S2) by plaquing ϕNM4γ4 on spacer F cells and sequencing the target to determine the escape mutations. The escaper was propagated on non-CRISPR staphylococci (*S. aureus* OS2) to high titers. Finally, 20,000 PFU of the escaper stock were added to the liquid HIA top agar mixture during the spacer acquisition assay described above.

### Determination of colony size over time

Images of the plates with ongoing spacer acquisition assays at 24 hours and 48 hours were acquired with a GE ImageQuant LAS 4000 Imager at a 1X magnification, and aligned using Adobe Illustrator.

### Microscopy analysis of colony morphology

Colonies on spacer acquisition plates were marked with pieces of glitter of different colors, which also served as a size marker. Images were taken with a 10X magnification with a Nikon SMZ-800N Stereo Zoom Microscope.

### Construction of pCRISPR(spacer F, Δcas2)

*cas2* was deleted from pCRISPR(spacer F) following the protocol used to convert pWJ40 into pRH61 [17].

### Treatment of colonies

0.5 μl of either 1X Phosphate Buffer Solution (PBS), 5 mM lysostaphin, or spacer F escaper phage stock were added on top of different colonies growing on plates.

### Time-shift assay

To measure the resistance of the bacteria to phage over time, top agar infection assays using founder F or C cells were performed as in Figures 1F or 1G, respectively. Samples were collected at 24, 36 and 48 hours post-infection by resuspending 12 of the resulting colonies in TSB, except for the experiment with founder F cells at 36 hours, for which 11 samples were recovered.

Phage from each sample was isolated and amplified as follows. First, 2 μL of each colony supernatant was spotted onto the surface of top agar seeded with non-immune *S. aureus* OS2 (for the phage used at time 0, we spotted 12 plaques obtained by plaquing the original stock of ϕNM4γ4 phage on top agar containing non-CRISPR hosts). Second, the phage “spots” obtained were resuspended into 20 μL of TSB and stored at 4°C.

To test the resistance of the bacteria over time, 2 μl of each colony resuspension were grown overnight in 200 μl of BHI media at 37°C. 150 μl of these cultures were mixed with 5 ml of HIA top agar containing 5 mM of CaCl_2_ and poured over plated on TSA plates. Time 0 bacteria were directly grown overnight at 37°C in BHI from our stocks of founder F and C cells. 2 μl of each amplified phage sample was spotted on top agar seeded with cells from different colonies (or founders in the case of time 0 infections). Plates were incubated overnight at 37°C and the resistance of the bacteria against the phage was scored as 0 (complete lysis), 0.5 (partial resistance), and 1 (full resistance).

## Supporting information

Supplementary sequences

## Acknowledgements

We would like to thank the Rockefeller University Genomics Resource Center for assistance with next generation sequencing experiments. L.A.M. is supported by a Burroughs Wellcome Fund PATH Award, and a NIH Director’s Pioneer Award (DP1GM128184). L.A.M. is an investigator of the Howard Hughes Medical Institute.

## Author contributions

N.C.P. and L.A.M. designed and conceived the study. N.C.P. performed all experiments. N.C.P. and L.A.M. wrote the manuscript.

---

**Figure S1.**
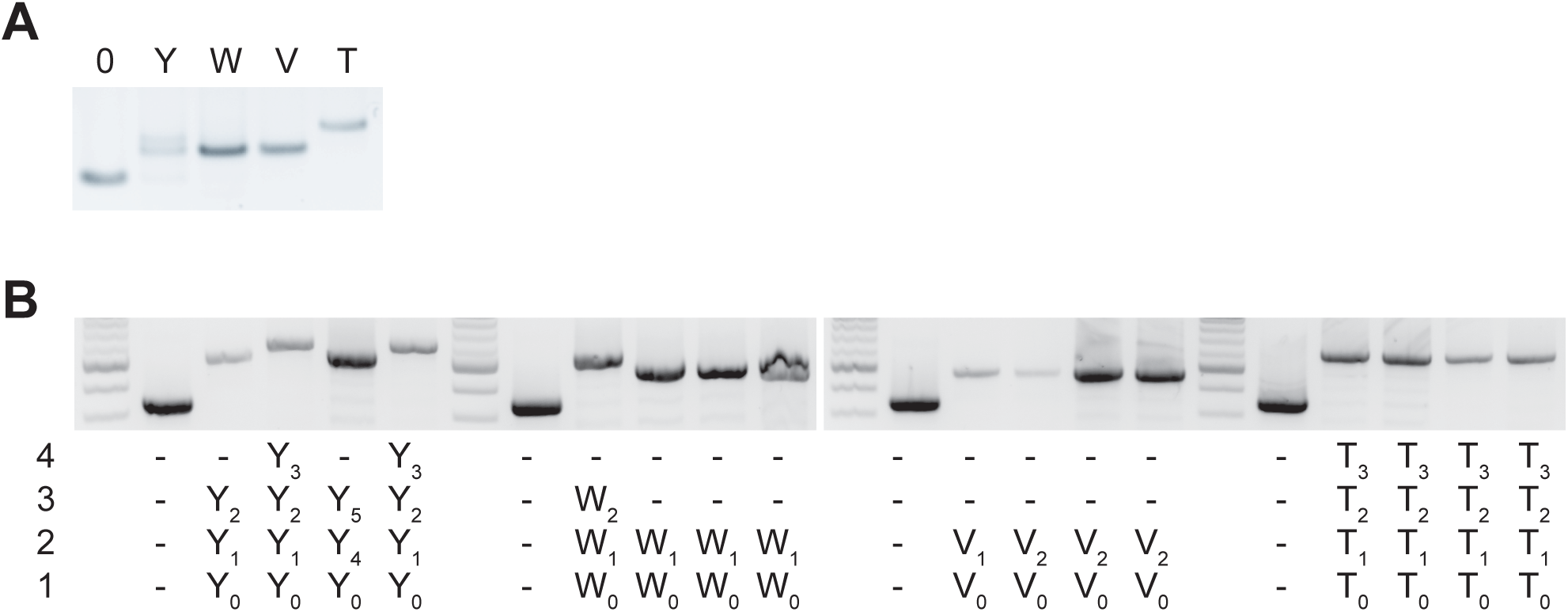
PCR analysis of the CRISPR array of re-streaked cells. (**A**) Agarose gel electrophoresis of the products of the amplification of the barcoded pCRISPR array present in staphylococci infected with ϕNM4γ4, using plasmid DNA templates extracted from bacteriophage-resistant colonies (labeled with different upper case letters) after infection in semi-solid (top agar) media, “0” indicates a no-spacer control sample. (**B**) Same as (**A**) but using template DNA extracted from colonies resulting from the re-streak of colonies shown in (**A**).

**Figure S2.**
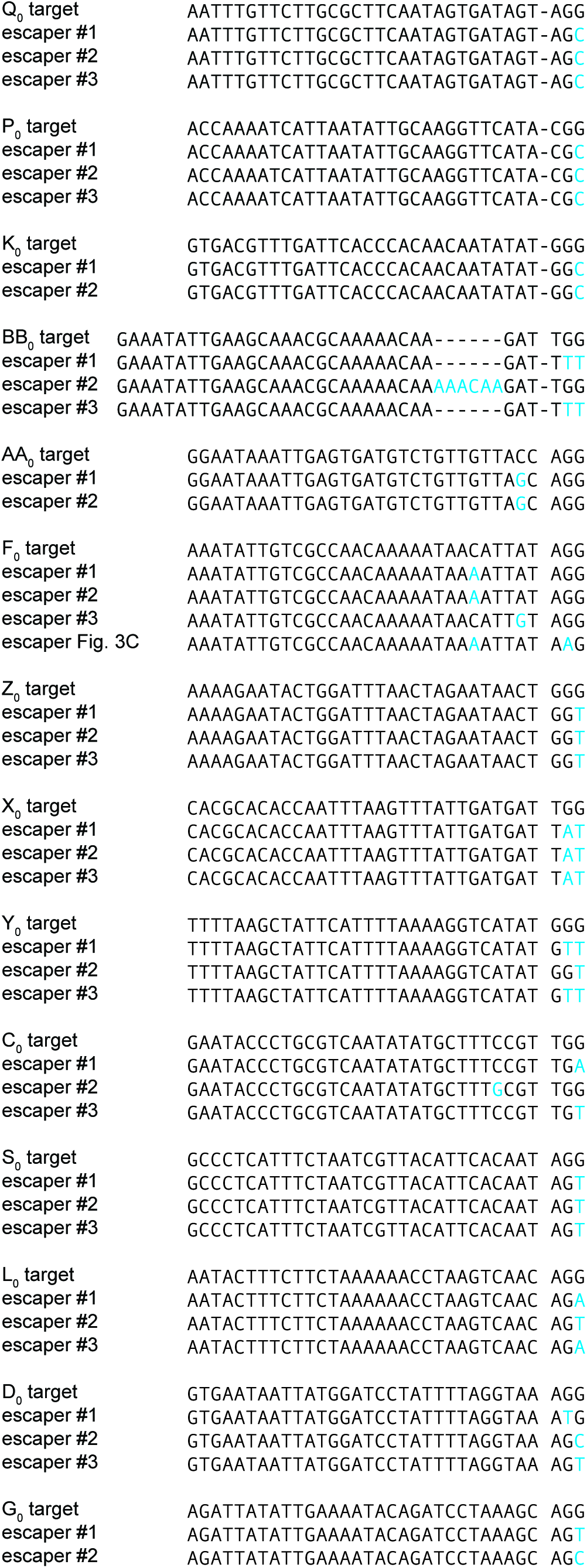
Target sequences of different escaper phages. DNA from 2-4 escaper plaques obtained in the experiment described in Figure 3A was isolated and the corresponding target region was amplified with target-specific primers and sequenced as described in Methods. Mutant sequences are aligned to the wild-type target (protospacer-PAM); mutations indicated in blue letters.

**Figure S3.**
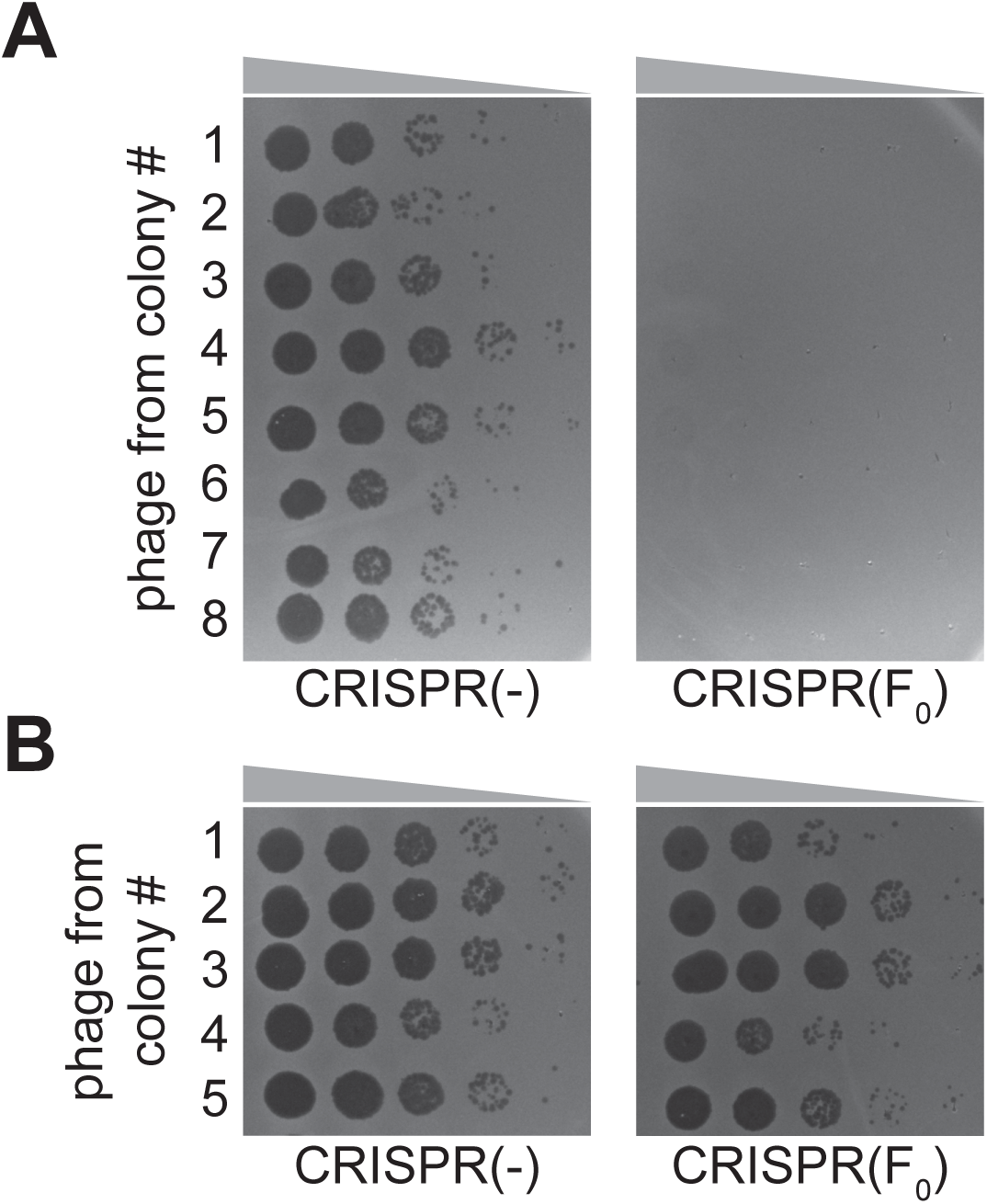
Quantification of escaper phage within spacer F founder colonies. (**A**) Phage was isolates from surviving colonies obtained in the experiment described by Figure 1F and 10-fold serial dilutions were spotted on lawns of sensitive staphylococci [CRISPR(-)], or staphylococci harboring the pCRISPR plasmid programmed with the founder spacer F [CRISPR(F_0_)]. (**B**) Same as (**A**) but using phage extracted from surviving colonies obtained in the experiment described by Figure 3C.

**Figure S4.**
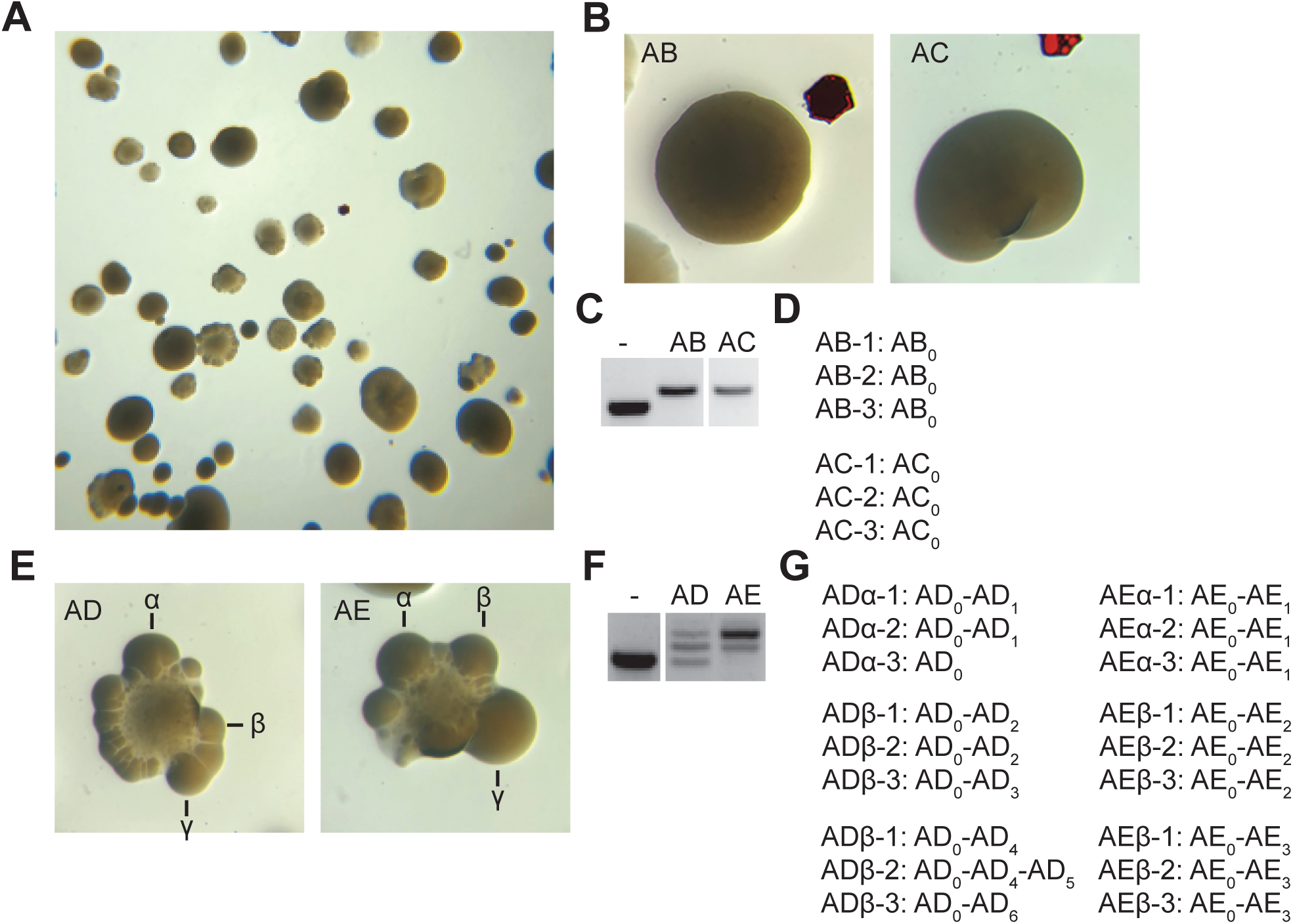
Analysis of additional smooth and sectored colonies. (**A**) Image of a plate containing staphylococci carrying pCRISPR that survive ϕNM4γ4 infection in solid media. (**B**) Image of two smooth colonies (containing the founder spacers AB and AC) from the plate shown in (**A**). (**C**) gel agarose analysis of PCR products obtained after amplification of the pCRISPR array of the smooth colonies shown in (**B**); (-) shows amplification of pCRISPR, a no-spacer control. (**D**) Sequencing results of the PCR products shown in (**C**). (**E**-**G**) Same as (**B**-**D**) but showing the analysis of two sectored colonies founded by spacers AD and AE; and also showing the sequencing of the pCRISPR array present in three different re-streaked sectors (α,β,γ).

**Figure S5.**
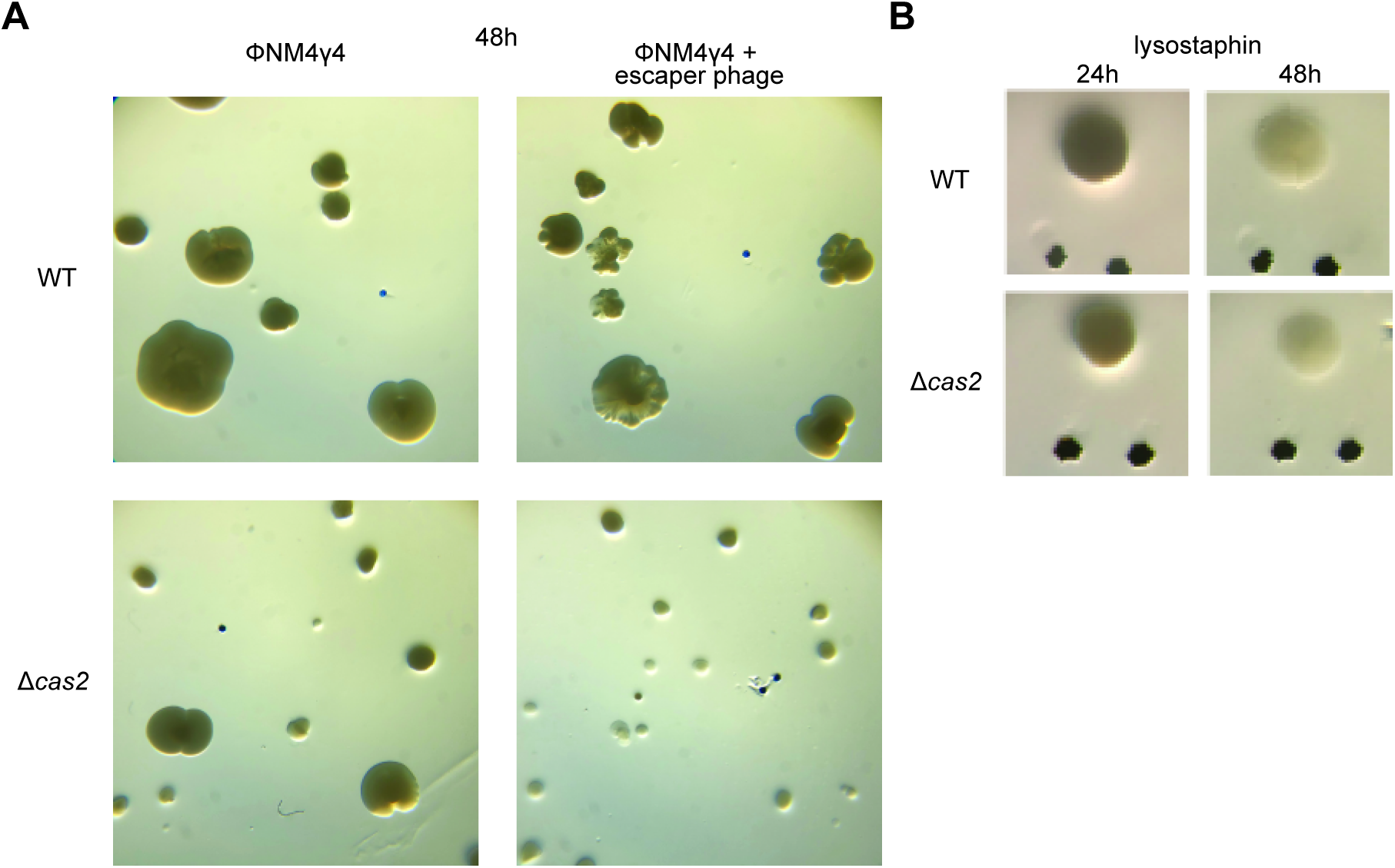
Requirement of spacer acquisition for the formation of sectored colonies. (**A**) Images of a plate grown for 48 hours after top agar infection of wild-type or Δ*cas2* mono-spacer founder F cells, with ϕNM4γ4 containing or not additional spacer F escaper phage. (**B**) Images of representative colonies of wild-type or Δ*cas2* mono-spacer founder F cells grown for 24 hours in the absence of phage, when a drop of the cell wall degrading enzyme lysostaphin was added on top and a second image was obtained 24 hours after. Glitter markers are shown to normalize both the position as well as the size of the image at different times.

